# Positive Selection and Adaptation of Novel Inner Ear Genes in the Mammalian Lineage

**DOI:** 10.1101/388496

**Authors:** Francisco Pisciottano, Alejandro R. Cinalli, Matías Stopiello, Valeria C. Castagna, Ana Belén Elgoyhen, Marcelo Rubinstein, María Eugenia Gómez-Casati, Lucía F. Franchini

## Abstract

The mammalian inner ear possesses functional and morphological innovations that contribute to its unique hearing capacities. The genetic bases underlying the evolution of this mammalian landmark are poorly understood. We propose that the emergence of morphological and functional innovations in the mammalian inner ear could have been driven by adaptive molecular evolution.

In this work, we analyzed the complete inner ear transcriptome in order to identify genes that show signatures of adaptive evolution in this lineage. We analyzed approximately 1,300 inner ear expressed genes and found that 13 % show signatures of positive selection in the mammalian lineage. Several of these genes are known to play an important function in the inner ear. In addition, we identified that a significant proportion of genes showing signatures of adaptive evolution in mammals have not been previously reported to participate in inner ear development and/or physiology. We focused our analysis in two of these novel genes: *STRIP2* and *ABLIM2* by generating null mutant mice and analyzed their auditory function. We found that mice lacking *Strip2* displayed a decrease in neural response amplitudes. In addition, we observed a reduction in the number of afferent synapses, suggesting a potential cochlear neuropathy.

Thus, this study shows the usefulness of pursuing a high-throughput evolutionary approach followed by functional studies to track down novel genes that are important for inner ear function. Moreover, this approach sheds light on the genetic basis underlying the evolution of the mammalian inner ear.

## BACKGROUND

The auditory system of mammals is characterized by a middle ear composed of three ossicles and an inner ear, with an auditory organ of Corti that possesses two types of specialized sensory hair cells (HCs), inner (IHCs) and outer hair cells (OHCs). The organ of Corti seems to have appeared early on in the evolution of therapsids (reptile ancestors of mammals) and it is derived from the basilar papilla of amniotes (1–3). In monotremes or egg-laying mammals (prototheria) the organ of Corti possess IHCs and OHCs but they are organized in several rows, whereas there is only one row of IHCs and three rows of OHCs in therian mammals (1–3), indicating further specialization in marsupials and placentals. Furthermore, the elongation of the organ of Corti contained in the cochlear duct in therian mammals (4) allowed the acquisition of unique hearing capacities among vertebrates, including high frequency sensitivity developed to an extreme level in some groups such as bats and whales (1, 5). In the mammalian cochlea IHCs and OHCs display a clear division of labor: IHCs receive and relay sound information behaving as the true sensory cells, while OHCs amplify sound information. Thus, IHCs which are the primary transducers, release glutamate to excite the sensory fibers of the cochlear nerve and OHCs act as biological motors to amplify the motion of the sensory epithelium. OHCs endowed mammals with a novel cochlear amplification mechanism known as somatic electromotility, crucial for auditory sensitivity and frequency selectivity (6, 7). In addition, these functional rearrangements at the inner ear level were accompanied by evolutionary changes at the auditory pathway in the central nervous system which underwent massive rearrangement in the tetrapod lineage (8).

The genetic remodeling that enabled the acquisition of morphological and functional novelties in the mammalian inner ear is poorly understood. Through mutant mice analysis, it has been shown that several transcription factors (TFs) are key to the development of the entire organ of Corti such as N-Myc, Pax2 and Gata3 (9–11). However, molecular evolutionary studies linking the evolution of these key developmental genes to the evolution of the mammalian inner ear are still missing. In this regard, recent findings have shown that several key inner ear genes underwent molecular adaptations that are probably linked to the evolution of the high-frequency hearing in mammals (12–14). For example, *SLC26A5* (encoding prestin), one of the first inner ear genes where signatures of positive selection in mammals were found (14), is a protein of the solute carrier (SLC) family directly involved in active amplification mechanisms of the mammalian cochlea (15, 16). Phylogenetic analyses of prestin and other SLC proteins have uncovered significant adaptive changes in the prestin amino-acid sequence, which have occurred essentially after the split between the avian and mammalian lineages. These adaptive changes have been related to the emergence of electromotility in mammals (12–14). In addition, we have recently shown that adaptive mutations of spectrin-βV occurred in the mammalian, but not the avian lineage, and were accompanied by substantial changes in the protein distribution within inner ear hair cells (17).

The deep evolutionary rearrangements that occurred in the mammalian inner ear involved the appearance of new cellular systems and novel functions, which probably required evolutionary changes in many proteins. In this work we aimed to identify the genetic bases underlying the evolution of the inner ear in mammals. To identify inner ear genes that adaptively evolved in the mammalian lineage we analyzed several expression databases and searched for signatures of positive selection by comparing synonymous and non-synonymous substitution rates in gene coding sequences. We found 167 inner ear expressed genes with signatures of positive selection in the mammalian lineage. Among them we discovered two previously unknown inner ear genes: *STRIP2* (from *Striatin Interacting Protein 2*) and *ABLIM2* (*Actin Binding LIM domain 2*) which were functionally characterized by generating novel strains of mutant mice by CRISPR/Cas9 technology. This work sheds light on the genetic basis underlying the evolution of the mammalian inner ear and highlights the usefulness of evolutionary studies to pinpoint novel key functional genes.

## Results

### Hair Cells Transcriptome Evolutionary Analysis

Several groups have recently sought to obtain the transcriptomes of cochlear hair cells, in attempts to reveal the genetic basis underlying the molecular mechanisms of hearing. Here, we analysed three mouse inner ear expression datasets generated by using different methodologies (18–20). To that end we developed a high throughput analysis pipeline that included an automatized method to retrieve and align the sequences of seven selected species representing all vertebrate classes and then run a branch-site evolutionary test (Figure 1, a). From these datasets we selected full-length protein coding genes presenting only one ortholog in each included species.

**Figure 1.**
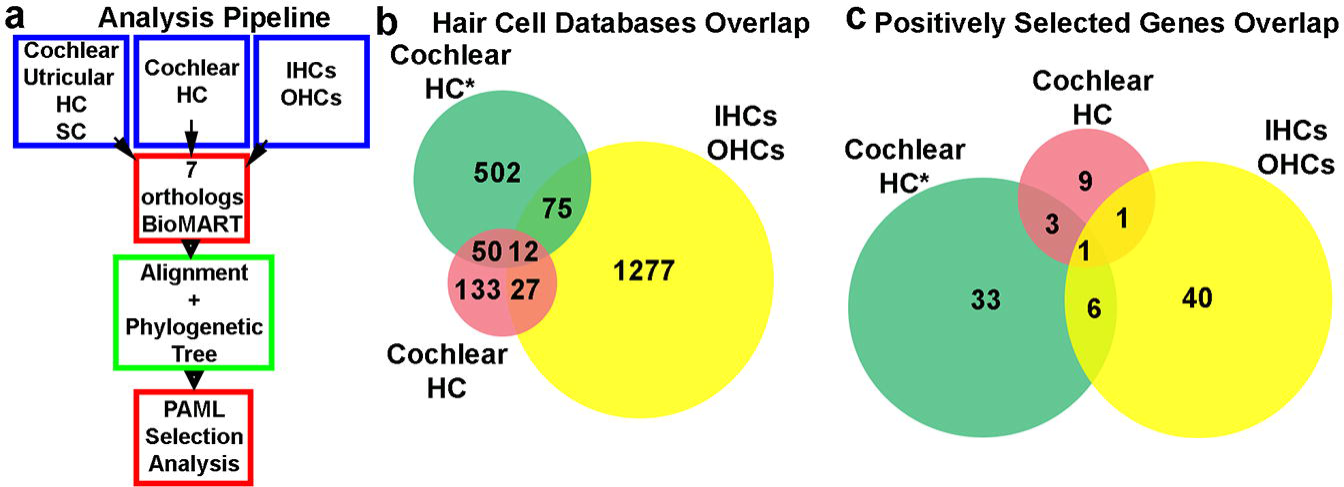
Analysis pipeline and different expression databases overlap. (a) An schematic diagram of the evolutionary analysis pipeline. (b) Gene overlap among the three cochlear hair cell expression datasets analysed. (c) Overlap among positively selected cochlear genes identified in the different databases. The only gene that is shared among the positively selected genes identified from the three databases is *STRIP*2.

We first analyzed transcripts differentially expressed in either IHCs or OHCs manually dissected from adult mouse cochleae (19). After filtering out the 1,193 reported IHC transcripts according to the aforementioned inclusion criteria we analysed 475 full-length coding transcripts by the branch-site method and detected 37 genes (8%) with signatures of positive selection in the mammalian lineage (Table 1). Then, we analyzed the reported 198 OHCs differentially expressed transcripts, of which only 88 fulfilled the inclusion criteria. The branch-site positive selection analysis for these 88 OHCs genes consistently retrieved 11 (12.5%) positively selected genes at p<0.05 (Table 1). Forty genes from these 198 OHCs transcripts were automatically excluded from our high-throughput pipeline due to one-to-many orthology relationships among the 7 selected species and 64 presented incomplete sequence information in one or more of the analysed species (Table S1). However, we decided to study the forty genes with complex orthology by unfolding the analysis in as many runs as necessary and found 4 additional positively selected genes (Table S2).

**Table 1.**
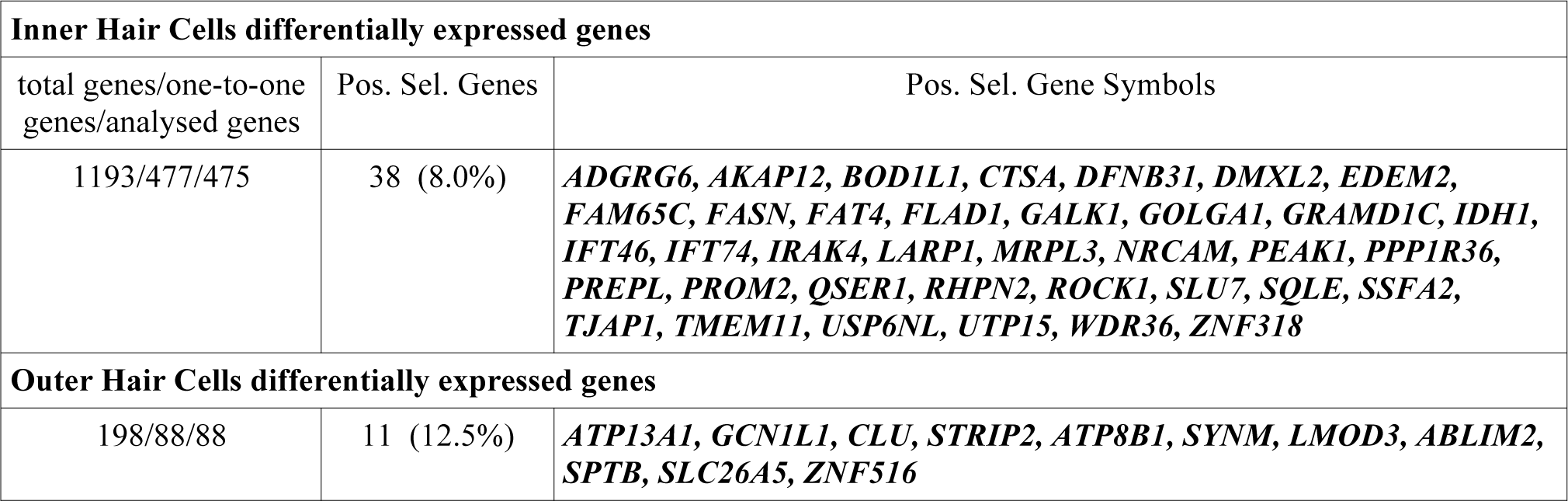
Positive selection analysis on differentially expressed IHCs and OHCs genes

The percentage of positive selected genes in OHCs (~12.5%, Table 1) was apparently greater than in IHC genes (8%, Table 1), and this seemed to be consistent with the greater morphological and functional reshaping that OHC underwent along the early evolutionary history of mammals. In contrast, as IHCs roughly maintained their ancestral role, it is possible that their ratio of positive selection was lower than the one we found in OHCs. However, a Fisher exact test showed that they are not significantly different (p= 0.30). This analysis indicated that both, IHCs and OHCs, went through similar levels of gene adaptive evolution probably underlying the morphological and functional remodelling that both cellular types underwent in the mammalian lineage.

The second dataset we analyzed used FACS isolation of hair cells from newborn mice (P0), which were then subjected to an RNA-Seq protocol (20). This study discriminated genes expressed in hair or surrounding cells form the cochlea and the utricle. From 82 postnatal cochlear hair cells enriched transcripts that met our analysis criteria, 14 (17.1%) showed to be under adaptive evolution in mammals (Table 2). From 16 postnatal utricular hair cell genes analyzed, we identified 4 (25%) showing signatures of positive selection in the basal mammalian tree branch (Table 2). In addition, from 390 genes expressed in inner ear surrounding or supporting cells we spotted 53 genes (13.6%; Table 2) with signatures of positive selection in mammals.

**Table 2.**
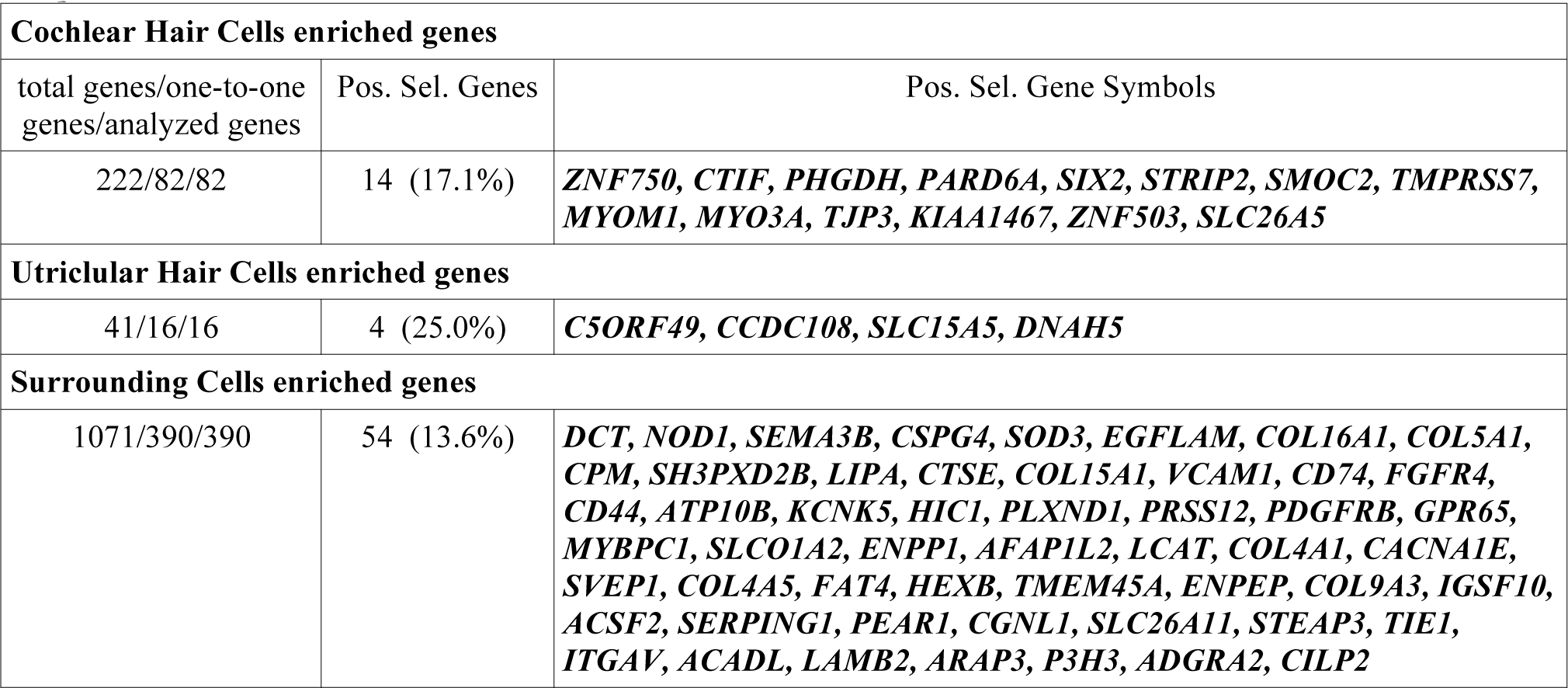
Positive selection analysis on genes enriched in hair and surrounding cells from cochlear and utricular samples

The third dataset analyzed was obtained by applying single-cell RNA-Seq transcriptomics to newborn mice inner ear cells. In this dataset we found 245 cochlear HC-specific genes that met our analysis criteria (18). Triplicate branch-site analysis consistently rendered the same 43 (17.5%) positive selected genes (p < 0.05; Table 3).

**Table 3.** Positive selection analysis on cochlear hair cells genes of newborn mice inner ears

| **Cochlear Hair Cells specific genes** |  |  |
| --- | --- | --- |
| total genes/one-to-one genes/analyzed genes | Pos. Sel. Genes | Pos. Selected Gene Symbols |
| 639/248/245 | 43 (17.5%) | ***CCDC189, TPRN, IQUB, DUSP27, LRP2, RHPN2, ABCC1, CFAP65, WDR78, GNMT, AP3B2, LAMC2, C9orf114, PGAP1, CDH23, PTPN6, DRC3, PEAK1, DGKG, ZZEF1, STRIP2, EFHC1, ADGRV1, SMOC2, DISP2, TMPRSS7, TJAP1, KCNS3, TJP3, GAS6, LMOD3, ABLIM2, OBSCN, MAP2, MMS19, DZIP1L, DMXL2, TMEM184A, CHRNA10, DLK2, SLC7A4, C2CD2L, KIF9*** |

In general, most of the genes that could not be analyzed by our approach were non-coding genes or presented incomplete orthology sequence information (one or more ortholog sequences were absent in genomic databases) or multiple orthology in one or more species (Table S1). The impossibility to analyze the evolution of these genes suggests that the number of genes that underwent positive selection in the mammalian lineage is possibly underestimated.

Since the databases we analyzed were mainly assembled using expression in postnatal stages and in order to better assess the evolutionary history of the mammalian inner ear, we also analyzed a manually collected dataset of transcription factors (TF) that have been reported to be key in inner ear development. Thus, we analyzed 23 TF and diffusible factors and found that most of them are extremely well conserved across vertebrates, suggesting that the evolution of their coding sequences did not play a significant role in mammalian inner ear changes. Only the gene helix-loop-helix 1 (*NHLH1)* exhibited signatures of positive selection in the mammalian lineage (Table S3).

### Overlap among different expression databases

When comparing the three cochlear hair cells datasets, we found that only 12 transcripts were present in all databases (Fig. 1, b). Even though the three studies aimed to obtain the HC transcriptome, the low level of overlap in the three datasets could be possibly explained by the different methodologies applied. While Scheffer and colleagues (20) isolated genes preferentially expressed in cochlear or utricular HC over those expressed in surrounding cells, Liu and collaborators (19) reported OHC genes differentially expressed in OHC in comparison to IHCs, and vice versa. Similar to Scheffer et al, Burns and colleagues (18), reported genes differentially expressed in HC in comparison to supporting cells, but only 62 genes were shared by these two datasets (Fig. 1, b). This low level of overlap suggests that the study of the three datasets ensured a more comprehensive analysis and includes a higher number of transcripts, increasing the chances of detecting more inner ear genes under positive selection.

Regarding the positive selected genes identified by the automatic pipeline, only *STRIP2* was present in the three databases (Figure 1, c). In addition, among 14 postnatal HC (18) and 49 IHC or OHC (11 from OHC and 38 from IHC) positively selected genes, we found only two genes present in both datasets: *SLC26A5* and *STRIP2* (Figure 1, c). On the other hand, between the 43 and 49 positively selected genes identified in the Burns et al (18) and the Liu et al. (19) datasets we found that 7 overlapped, *STRIP2, ABLIM2*, *Leimodin 3* (*LMOD3*)*, DMX like 2* (*DMXL2*)*, Rhophilin Rho GTPase Binding Protein 2* (*RHPN2*)*, Pseudopodium Enriched Atypical Kinase 1* (*PEAK1*) and *Tight Junction Associated Protein 1* (*TJAP1*) (Figure 1, c). Altogether, these data indicated that despite the source of the transcriptome and the small overlap among the databases a few genes are consistently present and identified as being under positive selection by our evolutionary analysis.

### Functional enrichment analysis of OHC and IHC differentially expressed genes

In order to analyze if the OHCs and IHCs positively selected genes shared associated functions we carried out a functional enrichment analysis using the DAVID tool for functional annotation (https://david.ncifcrf.gov/) (21). The results indicated that the most enriched functional term for the category of molecular function in the 11 OHCs positively selected genes was “cytoskeletal protein binding” (GO: 0008092) (p = 3.02e-5). Moreover, the second most enriched term of this category was “structural constituent of the cytoskeleton” (GO: 0005200) (p = 1.5e-2). The remaining seven enriched terms reported in the results were associated with transport through the membrane of ions and other molecules (Table S4). The enrichment in cytoskeleton functional proteins found in the 11 OHCs positive selected genes is specific for this particular gene subset and not product of an inherited overrepresentation of proteins related to cytoskeleton in the list of the 88 OHCs, since the GO terms that better apply to the genes in this dataset are from completely different functional categories (Table S4). We also performed a functional enrichment of the 11 positively selected genes, but this time using the list of 88 OHC genes analyzed, as background. This analysis returned a single functional enrichment term from these two lists, “cytoskeleton binding protein” (GO: 0008092) (2.2e-3), thus confirming the previous result (Table S5). A functional enrichment analysis was also performed for the 37 positively selected genes identified in the IHC differentially expressed genes dataset (19) and it did not yield any enriched functional GO term.

Altogether, these findings indicate that the OHC genes that underwent positive selection could have contributed to the acquisition of the highly specialized cytoskeleton present in these cells that underlies its distinctive functional properties, including somatic electromotility.

### Extended evolutionary analyses of OHCs positively selected genes

Because of the importance of OHC in mammalian audition, and to confirm the results of our exploratory analysis, we increased the statistical power of the evolutionary analysis by including more vertebrate species for all OHC genes displaying signatures of adaptive evolution (Table 1). For this analysis, we used all available sequences of vertebrate species and confirmed the results previously obtained (Table 4), demonstrating that our approach using seven representative vertebrate species is sufficiently robust to identify genes with strong signals of adaptive evolution.

**Table 4.**
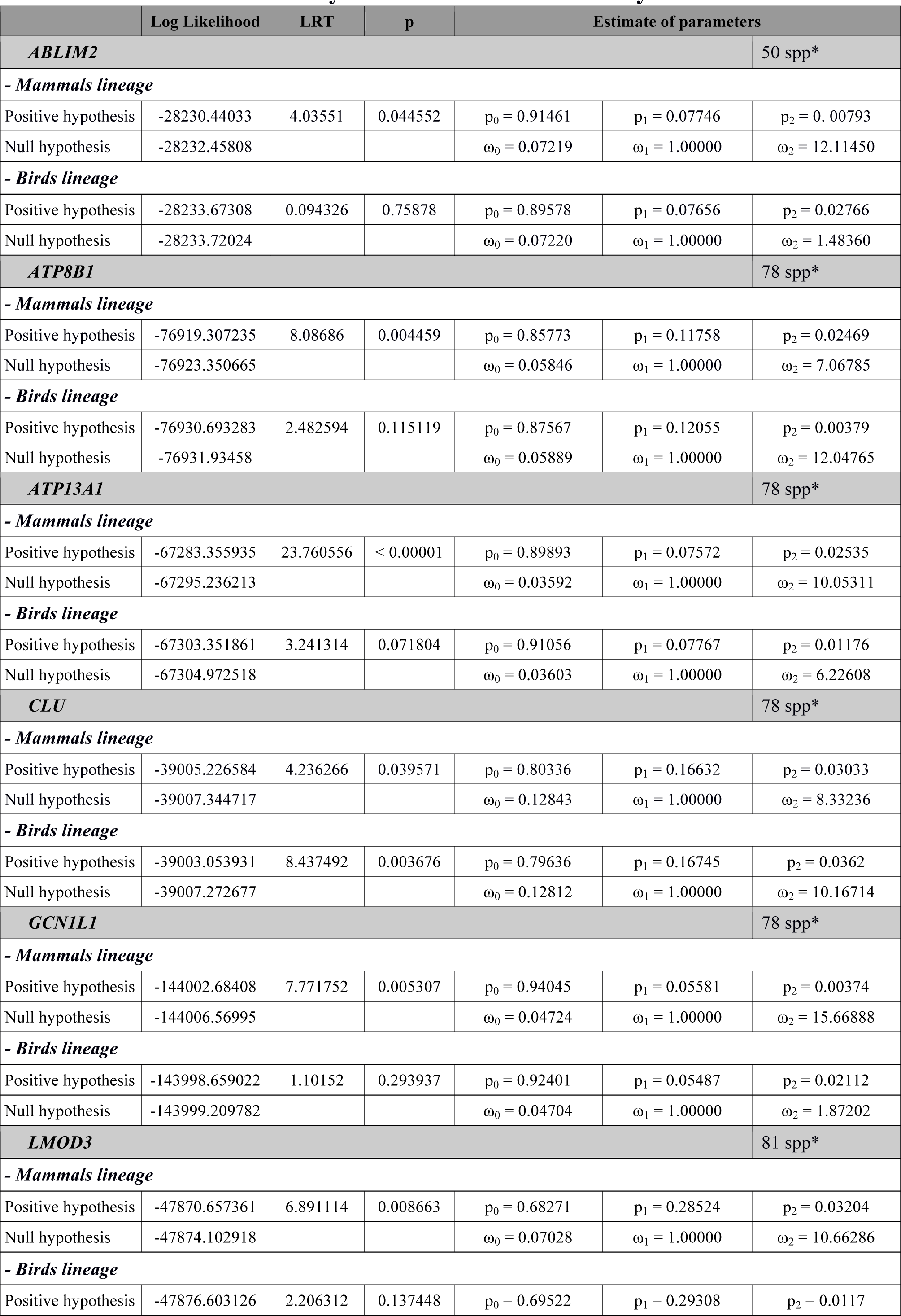
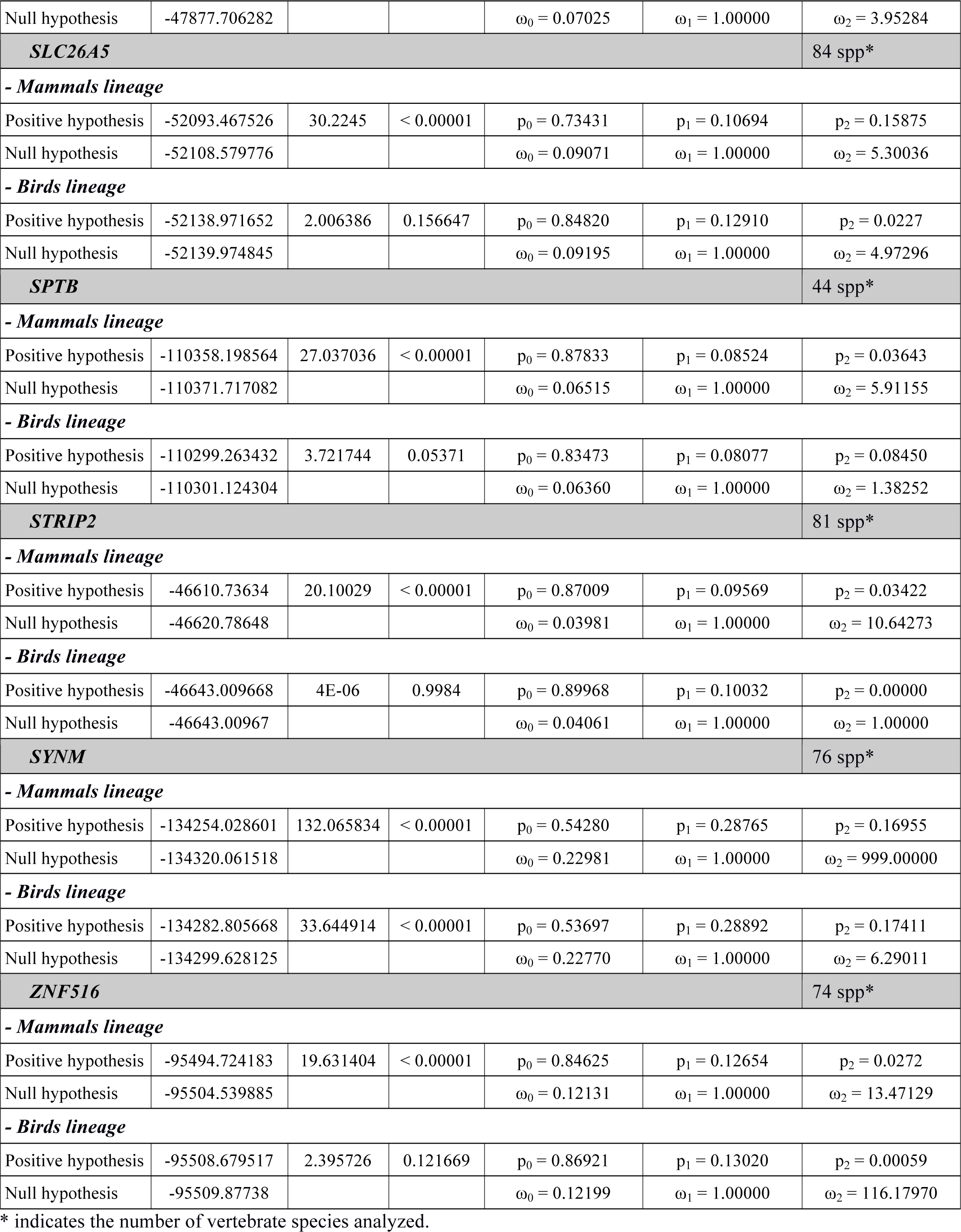
Outer Hair Cells Positively Selected Genes Extended Analysis

On the other hand, we tested the lineage leading to birds and found no evidence of positive selection for most genes except for synemin (*SYNM*) and clusterin (*CLU*) genes (Table S6).

We also tested the evolution of these OHCs genes in the tetrapod lineage and found signatures of positive selection, suggesting that remodelling of these genes happened in several moments of the vertebrate history (Table S6). Besides, in order to better assess the evolution of this lineage we analyzed the tetrapod lineage including the coelacanth *Latimeria menadoensis* sequence, a lobed-finned fish that posses a tetrapod-like ear but that lives in an aquatic environment. We found that in most genes the signature of positive selection is maintained when including this sequence (Table S6).

### Expression analysis of OHC positively selected genes

We next determined by in situ hybridization (ISH) and immunohistochemistry (IC) the expression pattern only of those OHC positively selected genes whose function in the inner ear is unknown. Thus, we did not include in this expression analysis *SLC26A5* and *Spectrin Beta I* (*SPTB*). We also excluded from this analysis the genes *ATPase Phospholipid Transporting 8B1* (*ATP8B1*) and *CLU.* It has already been shown that ATP8B1 is specifically located in the stereocilia of the inner ear HCs and that a mutated version causes hearing loss associated with progressive degeneration of the cochlear HCs (22). In addition, a recent detailed analysis of *CLU* showed that it is expressed in supporting cells of the inner ear and not in the HCs (23). Our IC assay shows that Lmod3 is strongly expressed in OHCs where it co-localizes with Prestin, whereas no expression was found in IHC or in supporting cells (Fig. 2, a, b, c). Our ISH results indicate that *Synm* and *General control of amino-acid synthesis 1 like 1* (*Gcn1l1*) genes are expressed in different cell types of the organ of Corti, including OHCs, IHCs and supporting cells; indicating that they are not HC-specific (Fig. 2, d, e, f, g, h, i). In contrast, ISH assays for *ATPase 13A1* (*Atp13a1*) (Fig. 2, j, k, l) and for *Zinc finger protein 516* (*Znf516*) (Fig. 2, m, n, o) failed to detect these genes to be expressed in the organ of Corti at the developmental stage analyzed (P8). In addition, our results indicated that *Strip2* is strongly expressed in OHCs, in IHCs and also in the spiral ganglion (Fig. 2, p, q, r, s). In addition, we found that *Ablim2* is expressed in both, OHCs and IHCs (Figure 2, t, u, v), and also in the spiral ganglion (Fig. 2, t, w). It is important to note that our expression analysis on positively selected genes in OHCs revealed some discrepancies with the original database where these genes were obtained. Liu *et al.* (19) examined the transcriptome of manually collected IHCs and OHCs from adult mouse cochleae and using microarray they reported 1,193 IHC and 198 OHC genes that were differentially expressed (p < 0.05) in one cell population over the other. Therefore, this database does not inform if these genes could be expressed in other cellular types of the inner ear. This explains why *Ablim2, Strip2, Gcn1l1* and *Synm* are also expressed in the spiral ganglion and in supporting cells, respectively.

**Figure 2.**
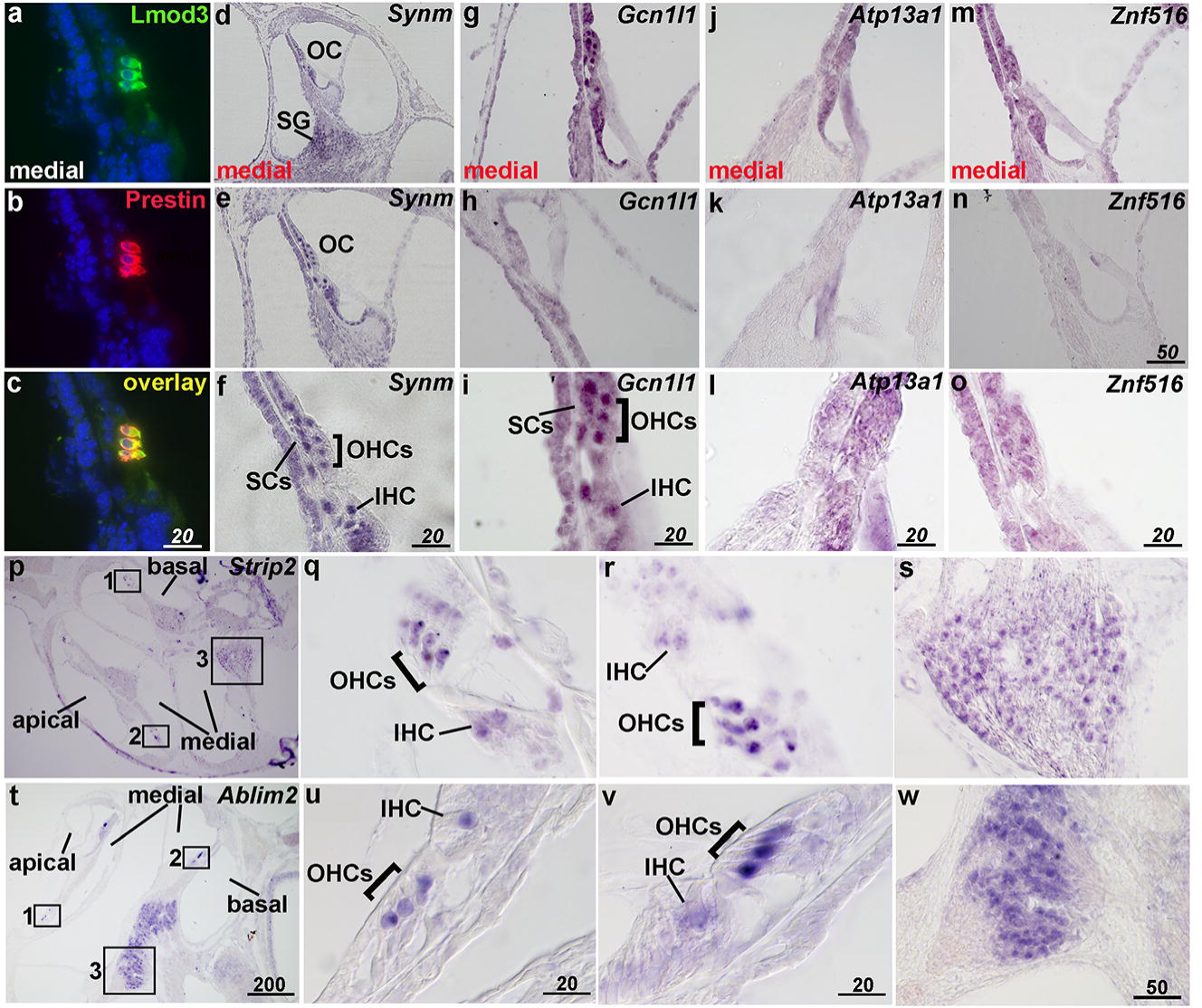
Expression analysis of OHCs positively selected genes in the inner ear. (a) Photomicrographs of IC assays showing Lmod3 expression (b) Prestin expression and (c) co-localization of these two proteins in OHCs. (d) In situ hybridization showing that *Synm* is expressed in the spiral ganglion and in other cellular types of the organ of Corti (e, f), including OHCs and IHCs. (g) *Gcn1l1* expression in the organ of Corti is localized in several cellular types. (h) Results obtained using the *Gcn1l1* sense probe. (i) A magnification of *Gcn1l1* expression showing IHC, OHC and SC in the organ of Corti. (j) *Atp13a1* expression in the organ of Corti obtained with the antisense probe (k) and with the sense probe. (l) A magnification of *Atp13a1* expression showing IHC, OHC and SC in the organ of Corti. (m) *Znf516* expression in the organ of Corti obtained with the antisense probe and (n) with the sense probe. (o) A magnification of *Znf516* expression showing IHC, OHC and SC in the organ of Corti. (p) Low magnification photomicrograph showing *Strip2* differential expression in OHCs and IHCs in the organ of Corti across different turns of the cochlea and in the spiral ganglion (q, r) Higher magnification of the section across the organ of Corti indicated by the squares 1 and 2 (s) Higher magnification of the spiral ganglion indicated by square 3. (t) Low magnification photomicrograph showing *Ablim2* differential expression in OHCs and IHCs in the organ of Corti across different turns of the cochlea and in the spiral ganglion (u, v) Higher magnification of the section across the organ of Corti indicated by the squares 1 and 2 (w) Higher magnification of the spiral ganglion indicated by square 3.

Altogether, these results allowed us to select those genes expressed in IHC and/or OHC, for follow-up in-depth evolutionary and functional studies. Taking into account our evolutionary and expression analysis results together, we decided to further study *STRIP2* and *ABLIM2* since their participation in inner ear function is unknown.

### Comprehensive evolutionary analysis and functional characterization of *STRIP2*

To better understand the impact of the evolutionary remodelling on *STRIP2* we identified particular amino acids displaying the signature of positive selection in mammals. To do that, after applying the branch-site positive selection test to the basal branch of the mammals (Fig. 3, a) we performed a follow-up Bayesian analysis that calculates the posterior probabilities that a site is under adaptive evolution. The results show a strong signal of adaptive molecular evolution in the mammalian lineage in 11 sites (Figure 3, b and c). Four of these positively selected amino acids were located in the N1221 domain, an acidic polypeptide with several possible transmembrane regions which function in still unknown (Figure 3, c).

**Figure 3.**
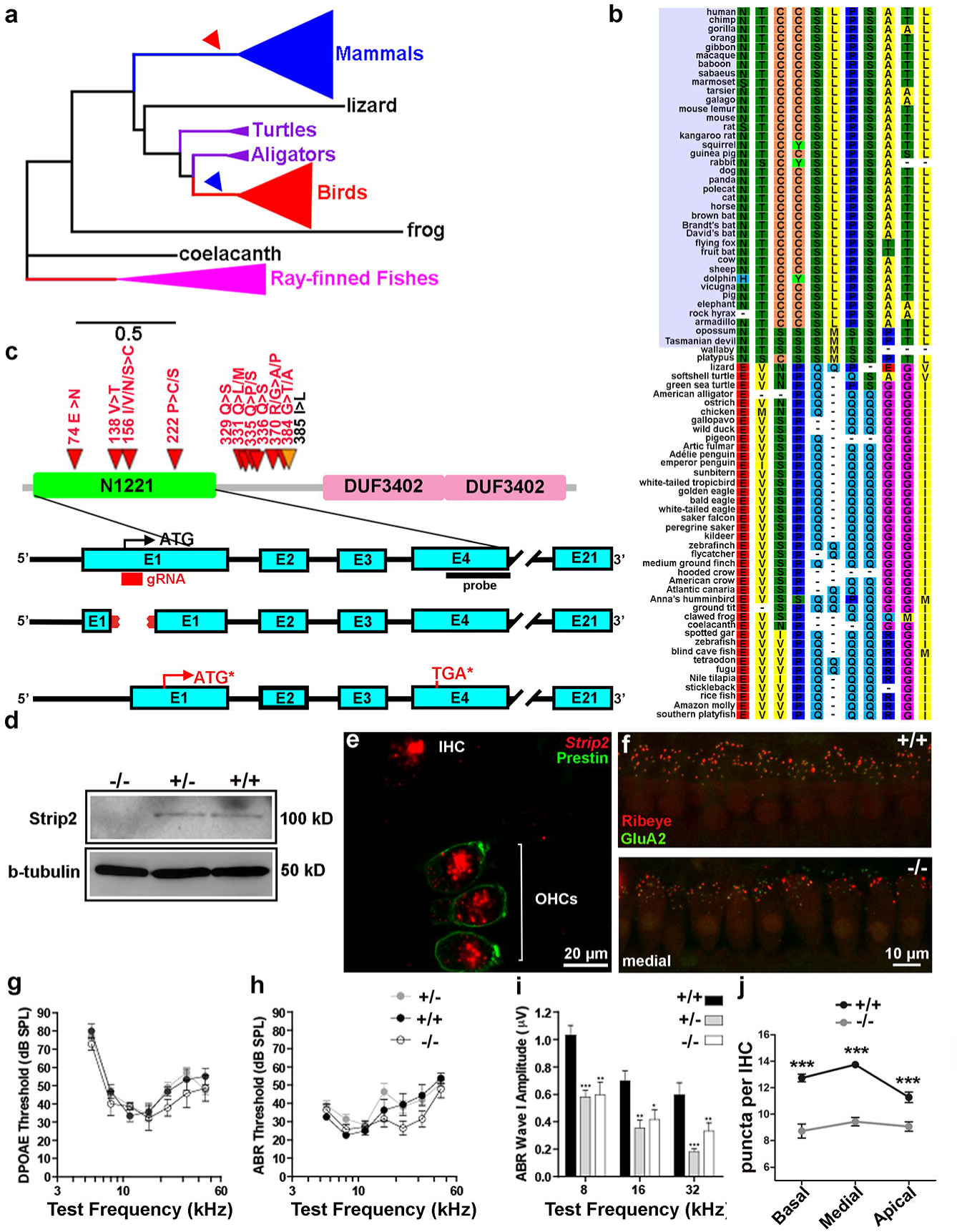
*STRIP2* is expressed in OHCs and IHCs, evolved adaptively in the mammalian lineage and it is important for IHCs function. (a) Phylogenetic tree of the 81 analyzed vertebrate species where some lineages are shown as collapsed branches for clarity. The mammalian and bird lineages (focal branches of the evolutionary analysis) are indicates with red and blue arrows, respectively. (b) Detail of the positively selected sites in mammals showing the changes in amino acids in the different vertebrates analyzed. (c) Top: Schematic drawing of the Strip2 protein showing the different functional domains. Red and yellow arrows indicate the location in the different functional domains of the positively selected sites in the mammalian lineage. Bottom: Scheme of the strategy developed to generate the mutant mice pedigree lacking Strip2. The sites of priming of the guide RNA on exon 1 and for the probe used for ISH on exon 4 are indicated. The deleted region affecting this exon is also shown. This deletion predicts the creation of an additional initiation codon (ATG*) a premature STOP codon (TGA*) in exon 4. (d) Western blot showing strong Strip2 expression in wild type mouse heart samples (*Strip2 ^+/+^*) a tissue where Strip2 is strongly expressed (24). We observed lack of expression in *Strip 2 ^-/-^*. (e) Photomicrographs of *in situ* hybridization and immunohistochemistry assays showing the expression of *Strip2* and Prestin in IHCs and OHCs in mouse cochleas (f) Representative confocal images of IHCs synapses from the medial turn of the cochleae immunolabeled for pre-synaptic ribbons (CtBP2-red) and post-synaptic receptor patches (GluA2-green) in *Strip 2 ^-/-^* and ^+/+^. Anti-CtBP2 antibody also weakly stains IHC nuclei. (g) DPOAEs and (h) ABRs thresholds measurements in *Strip 2 ^-/-^*, ^+/-^ and ^+/+^ in 2 months-old mice at different frequencies (From 5.6 to 45.25 kHz). (i) ABR wave I amplitude at 8, 16 and 32 kHz. Statistical analysis: one-way ANOVA followed by Tukey’s Multiple Comparison Test; * = p<0.05; ** = p<0.01; *** = p<0.001. (j) Synapsis quantification and statistical analysis.

To generate *Strip2* mutant mice we used a single guide RNA directed to exon 1 that induced a 64 bp deletion, which predicts the generation of an inactive truncated protein (Fig. 3, c). We verified the generation of *Strip2* mutations in the engineered mice by genomic DNA sequencing and also by Western blot protein analysis (Fig. 3, d). We observed a lack of Strip2 protein in the hearts of Strip2 ^-/-^ mice (Fig. 3, d), a tissue where Strip2 is strongly expressed (24).

We performed auditory functional studies of *Strip2* newly generated mutant mice by means of two complementary techniques that allow differential diagnosis of OHC versus IHC/neuronal dysfunction throughout the cochlea. To evaluate the integrity of the hearing system we recorded ABRs (Auditory Brainstem Responses) that are sound-evoked potentials generated by neuronal circuits in the ascending auditory pathways (25). We also evaluated the OHCs function through distortion product otoacoustic emissions (DPOAE) testing (26).

We measured DPOAEs thresholds at different frequencies (from 5.6 to 45.25 kHz) in *Strip2* ^-/-^, ^+/-^ and ^+/+^ littermates at 2 months. Remarkably, there were no changes in DPOAEs thresholds in the three genotypes (*Strip2* ^+/+^ n=8; *Strip2* ^+/-^ n=18; *Strip2* ^-/-^ n=7; Fig. 3, g). These results show that the lack of *Strip2* does not affect the OHCs’ biological motors. In addition, we found no differences in ABR thresholds at all the frequencies tested (*Strip2* ^+/+^ n=8; *Strip2* ^+/-^ n=18; *Strip2* ^-/-^ n=7; Fig. 3, h). However, when analyzing ABR wave I, we found a large reduction in amplitudes at 8, 16 and 32 kHz in both *Strip2* ^-/-^ and ^+/-^ mice (*Strip2* ^+/+^ n=8; *Strip2* ^+/-^ n=18; *Strip2* ^-/-^ n=7; Figure 3, i). These results might indicate a potential cochlear neuropathy. The ABR wave I latencies were not altered in the different genotypes, suggesting that the lack of *Strip2* does not affect cochlear nerve conduction. Through ISH analysis we verified that *Strip2* is expressed at the cochlear nucleus (CN; Fig. S1). However, since no alteration in wave II ABRs amplitudes were observed, no overt malfunction of synaptic transmission at this central relay is expected.

To analyze if the reduction in ABR peak I amplitude is due to the loss of auditory nerve synapses we immunostained whole-mount organs of Corti with antibodies against CtBP2-Ribeye, a critical protein present at the presynaptic ribbon (27), and GluA2 AMPA-type glutamate receptors, which are expressed at the postsynaptic afferent terminal (28). In each immunostained whole mount organ of Corti we counted the colocalized synaptic puncta at the medial, apical and basal cochlear regions. We found that the number of synaptic puncta per IHC was significantly reduced in the three regions analyzed (basal *Strip2^+/+^* 12.73 ± 0,2944 and *Strip2*^-/-^ 8.729 ± 0.5265 t=6.251 df=233 p<0.001; medial *Strip2*^*+/+*^ 13.74 ± 0.2209 *Strip2*^-/-^ t=10.37 df=278 p<0.001; apical *Strip2*^*+/+*^ 11.27 ± 0.3958, *Strip2*^-/-^ 9.066 ± 0.3580 t=4.095 df=157 p<0.001; n = 5 for both genotypes; mean ± SEM; Figure 3, f and j).

### Comprehensive evolutionary analysis and functional characterization of *ABLIM2*

In order to identify amino acids sites displaying the signature of positive selection we performed an exhaustive evolutionary analysis using 50 *ABLIM2* vertebrate sequences (Fig. 4, a and c). Our analysis, identified two amino acids (518 L/P >S; 557 E/A>N) that displayed signatures of positive selection, both located in the Adherens-junction domain (Fig. 4, b and c).

**Figure 4.**
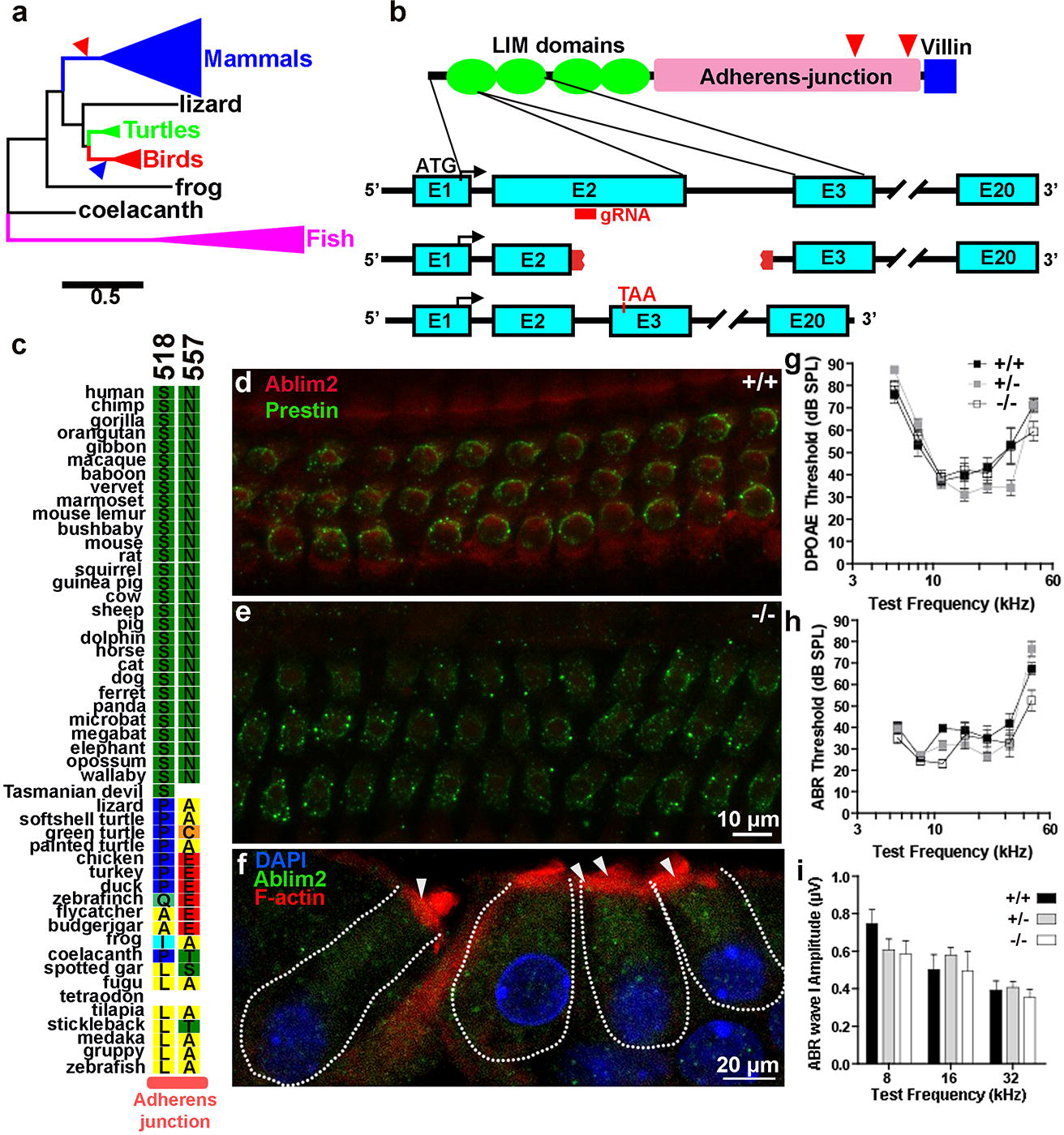
*ABLIM2* is expressed in OHCs and IHCs and evolved adaptively in the mammalian lineage. (a) Phylogenetic tree of the 50 analyzed vertebrate species where some lineages are shown as collapsed branches for clarity. The mammalian and bird lineages (focal branches of the evolutionary analysis) are indicated with red and blue arrows, respectively. (b) Top: Schematic drawing of the ABLIM2 protein showing the different functional domains. Red arrows indicate the location in the Adherens-junction domain of the positively selected sites in the mammalian lineage. Bottom: Scheme of the strategy developed to generate the mutant mice pedigree lacking Ablim2. The site of priming of the guide RNA on exon 2 is indicated. The deleted region affecting this exon is also shown. This deletion is predicted to generate a premature STOP codon in exon 3. (c) Detail of the amino acids positively selected in the different vertebrates analyzed. (d, e) Immunohistochemistry assays showing the expression of Ablim2 and Prestin in *Ablim2 ^-/-^ and ^+/+^* representative mice. (f) Immunohistochemistry showing expression of Ablim2, F-actin and DAPI in OHCs and IHCs. White arrows indicate colocalizton spots in the stereocilia (g) DPOAEs (top) and (h) ABRs (bottom) thresholds measurements in *Ablim2 ^-/-^, ^+/-^ and ^+/+^* in 2 months-old mice at different frequencies (From 5.6 to 45.25 kHz). (i) ABR wave I amplitude at 8, 16 and 32 kHz. Statistical analysis: one-way ANOVA followed by Tukey’s Multiple Comparison Test.

Since it has been reported that *Ablim2* interacts with F-actin (29) we analyzed the evolution of genes encoding actin in the inner ear (*ACTB* and *ACTG; (30)*) in the mammalian lineage. We found no signatures of positive selection in any of these genes in the lineage leading to mammals (p=1;Table S7).

In addition, to assess if F-actin and Ablim2 co-localized in hair cells of the mammalian cochlea we performed immunostaining of both proteins. Our results indicated that Ablim2 is mainly distributed in the cytoplasm and that both proteins co-localized at the stereocilia of inner and outer hair cells (Fig. 4, f).

Mutant mice lacking *Ablim2* were generated by deleting 202 base pairs comprising the last 48 aminoacids of exon 2 and 58 bp of the second intron (Fig. 4, b). Immunohistochemical analysis confirmed that Ablim2 is absent in the inner ear of *Ablim2* ^-/-^ mice (Fig. 4, d, e). We analyzed the auditory function in mice lacking *Ablim2* and found that cochlear thresholds were not affected when measured by DPOAEs or ABRs (*Ablim2* ^+/+^ n=11; *Ablim2* ^+/-^ n=18; *Ablim2* ^-/-^ n=8; Fig. 4, g, h). The earliest wave of the ABR (wave I), which represents the summed sound-evoked spike activity at the first synapse between IHCs and afferent nerve fibers, was not modified in amplitude or latency in *Ablim2* ^-/-^ mice at 8, 16 or 32 kHz (*Ablim2* ^+/+^ n=11; *Ablim2* ^+/-^ n=18; *Ablim2* ^-/-^ n=8; Fig. 4, i). These results suggest that the absence of *Ablim2* does not affect either cochlear amplification or auditory nerve function.

## Discussion

To unravel the evolutionary history of the mammalian auditory system we performed a detailed evolutionary analysis of several databases reporting genes expressed in the inner ear. We analyzed approximately 1,300 coding gene sequences and identified 167 displaying signatures of positive selection in the mammalian lineage. Since we are particularly interested in the evolution of the cochlea in mammals we focused our analysis in genes expressed in OHCs and IHCs. We surprisingly found that a similar proportion of IHC and OHC genes underwent positive selection, suggesting that both cellular types underwent significant functional remodeling in the mammalian lineage. We selected two of these genes, *ABLIM2* and *STRIP2*, to perform functional studies that included the generation of mutant mouse strains lacking these proteins. We discovered that *Strip2 ^-/-^* and *Strip2^+/-^* mice exhibited a large reduction in ABR peak I amplitudes, suggesting that *Strip2* plays a functional role in the first synapse between IHCs and afferent nerve fibers. Moreover, cochlear sensory epithelium of *Strip2 ^-/-^* immunostained for pre- and post-synaptic markers showed a reduction in auditory-nerve synapses, suggesting that lack of *Strip2* expression leads to cochlear synaptopathy.

The evolutionary analysis of OHCs genes identified 11 genes showing strong signatures of positive selection in the mammalian lineage. This might aid to unravell the evolution of the genetic network that shaped the remarkable functional motor capacities of this mammalian cell type. These include genes such as *SLC26A5* (encoding prestin) and *SPTB* (erythrocytic spectrin beta 1) previously identified as key players in the evolution of the mammalian inner ear, which underwent selective pressure in mammals and are expressed in OHCs (12-14, 17).

Within the 198 OHCs differentially expressed genes, some particular genes previously shown to be strongly expressed in OHCs and also to have undergone positive selection in the mammalian lineage were absent. These include the nicotinic acetylcholine receptor subunit alpha 10 (*Chrna10*), and Spectrin Beta, Non-Erythrocytic 5 (*Sptbn5*) (13, 14, 17). Although *Chrna10* was reported as the most differentially expressed gene in OHCs (19), this gene was filtered out in our automatic pipeline because it has been duplicated in the fish lineage (14). However, we have previously carried out deep evolutionary studies with this nicotinic receptor and shown that it underwent positive selection and functional remodeling in the mammalian lineage (13, 14, 31). On the other hand, since *Sptbn5* is expressed at similar levels in both, IHCs and OHCs, it was not found as differentially expressed in one cell type over the other (19). To our surprise*, Sptbn5* was not reported in the other two HC expression databases analyzed in this study (18, 20).

Our data highlight that several OHC genes underwent positive selection probably underlying some of the morphological and functional changes that this cellular type experienced throughout mammalian evolution. In addition, it is interesting to note that we surprisingly found a similar proportion of genes expressed in IHCs that went through adaptive evolution in the mammalian lineage. Moreover, the present expression analysis shows that most of the genes previously reported to be differentially expressed in OHCs (Liu et al., 2015), are in fact strongly expressed in IHCs and other cell types in the mammalian inner ear. Thus, we show that IHCs and OHCs underwent similar levels of evolutionary driven genetic remodeling. This might underlie the profound division of labor that IHCs and OHCs play in the mammalian cochlea: behaving as true sensory cells in the case of IHCs and performing cochlear amplification in the case of OHCs. Division of labor has also been observed in hair cells of the avian auditory papilla (the equivalent to the cochlea in mammals), which possess tall (THCs) and short hair cells (SHCs) (32). However, current data indicate that even though SHCs and THCs in the chicken auditory papilla show analogies with OHCs and IHCs, they differentiate in several aspects: prestin is highly expressed in both cellular types, and voltage-induced bundle motion is not confined to the SHCs since it is also observed in THCs (33). Thus, the mammalian specific division of labor between IHCs and OHCs is consistent with the extensive molecular adaptations that we have uncovered and that most probably underlie the functional specialization of these mammalian phenotypical novelties. Thus, our data provides a new framework for analyzing the evolution of the mammalian inner ear in which parallel evolution of IHCs and OHCs occurred in order to accommodate the specific and exclusive functions of each cellular type.

Taking into account our evolutionary findings and the gene expression study outcome, we selected *ABLIM2* and *STRIP2* to perform a comprehensive assessment of positive selection in vertebrates and functional analysis through the generation of null mutant mice where we examined auditory function. Our results indicated that both, *ABLIM2* and *STRIP2* displayed signatures of adaptive evolution in mammals but not in the lineage leading to birds (Aves), suggesting that other proteins were responsible for the evolutionary changes observed in the inner ear in this group of vertebrates. In addition, both *ABLIM2* and *STRIP2* evolved adaptively in the tetrapod lineage suggesting that they might also be important for the remodeling of the inner ear that led to the appearance of the basilar papilla, the precursor to the organ of Corti (for a review see (1)).

Our functional studies indicated that *Strip2* (also known as *FAM40B* or Myoscape), is important in inner ear physiology since mice lacking *Strip2* show an impaired first synapse between IHCs and afferent nerve fibers as evidenced by a reduction of the number of afferent synaptic contacts and of the ABR peak I amplitude. It has been recently shown that Strip2 directly interacts with the C-terminal tail domain of the L-type Ca2+ channel in cardiomyocytes (24). Knockdown of *Strip2* in cardiomyocytes results in downregulation of Ca2+ channel expression together with a reduction of calcium channel currents (24). In the inner ear, hair cell neurotransmission is thought to mainly rely on Ca_V_1.3 L-type Ca2+ channels (34–38). In addition, it has been proposed that L-type Ca2+ channels may have a role in hair cell development. Before the onset of hearing in altricial rodents, IHCs fire Ca2+ action potentials (39–41), which drive afferent synaptic transmission (39–42). This presensory activity is probably important for the development and maintenance of the auditory pathway (43, 44). Mice lacking Ca_V_1.3 channels are deaf and finally undergo degeneration of afferent auditory nerve fibers and hair cells (45). We can hypothesize that, in the inner ear, *Strip2* may interact with Ca_V_1.3 channels promoting the correct assembling and/or the maintenance of auditory nerve synapses. It would be interesting to investigate if at the onset of hearing the proper number of synapses per IHCs is formed and degenerate afterwards or if they never developed. Nonetheless, our results show that despite the loss of auditory nerve synapses, ABRs thresholds were not elevated, a clear indication of cochlear neuropathy (46), indicating that *Strip2* might be important for the survival of low-spontaneous rate auditory nerve fibers.

It is interesting to note that *Strip2* plays a role in cell shape and cytoskeleton organization (47) and that the functional properties of OHCs are largely based in a highly specialized cytoskeleton linked to the cytoplasmic membrane that allows their remarkable conformational changes (6). Moreover, our functional annotation analysis indicates that genes linked to cytoskeleton functions are over-represented among OHCs positively selected genes. The fact that DPOAEs thresholds were normal in *Strip2* ^-/-^ mice indicates that the lack of this protein does not affect the OHCs’ electromotile function.

On the other hand, our functional studies indicated that mice lacking *Ablim2* display normal cochlear thresholds, as measured by both, DPOAEs and ABRs. These results might suggest that Ablim2 does not participate in cochlear amplification or in IHCs function or that other functionally redundant gene could compensate its absence. Our expression pattern analysis of *Ablim2* in the inner ear indicates that it is highly expressed in OHCs and IHCs of the organ of Corti, and also in the spiral ganglion. *ABLIM2* is a STARS (striated muscle activator of Rho signalling) protein that strongly binds to F-actin (29). It has been shown that *Ablim2* is highly expressed in muscle where, it apparently regulates the sensing of biomechanical stress and also in the brain where it participates in axon guidance (29). Our colocalization studies indicate that Ablim2 is expressed in the cytoplasm of hair cells and it seems to colocalize with F-actin at the stereocilia. Since *ACTB* and *ACTG,* actin-encoding genes, did not evidence signatures of positive selection, they did not undergo parallel evolution with *ABLIM2*. Further functional studies will be necessary to determine the functional relevance of this protein in inner ear physiology.

It is interesting to analyze our results in the light of one of the most fundamental debates in adaptive evolution: does evolution proceed primarily through changes in protein-coding DNA or in noncoding regulatory sequences? It has been hypothesized that morphological adaptation occurs mainly via noncoding changes, since many developmental genes are active in many tissues, whereas noncoding mutations can modify a gene’s activity in just one or two tissues, avoiding pleiotropic effects (48). In contrast, it has been also suggested that adaptive evolution of protein-coding sequences, as well as other mechanisms like gene duplication, play a significant role in the evolution of form and function (49). In addition a combination of these two mechanisms seems to also play a role through the duplication of genes followed by subfunctionalization (i.e. duplicated genes or paralogs, retain a subset of its original ancestral function or acquire different expression domains) (50, 51). In this regard, the co-option of new regulatory machinery seems to be the mechanism that facilitates the acquisition of novel expression domains (for a review see (52). In the evolution of the inner ear, the several steps that led to the appearance of the complex sensory organ of vertebrates (53) must have probably involved all these evolutionary mechanism actin on many genes and gene regulatory networks. Therefore, more comparative analysis about the evolution of protein coding and noncoding sequences that control the expression of genes in the developing and adult inner ear would be necessary to better illuminate our understanding of functional and morphological evolution of this mammalian sensory organ.

In summary, through this evolutionary approach we discovered that *STRIP2* underwent strong positive selection in the mammalian lineage and plays an important role in the physiology of the inner ear. Moreover, our combined evolutionary and functional studies allow us to speculate that the extensive evolutionary remodeling that this gene underwent in the mammalian lineage provided an adaptive value. Thus, our study is a proof of concept that evolutionary approaches paired with functional studies could be a useful tool to uncover new key players in the function of organs and tissues.

## Methods

### Sequences Retrieval

Selected species for this study were chosen in order to maximize the presence of reported orthologs for the majority of genes analyzed, taking care to keep a balance between mammalian and non-mammalian vertebrate species. Our default chosen species for this study were: human (*Homo sapiens*), mouse (*Mus musculus*), dog (*Canis familiaris*), opossum (*Monodelphis domestica*), chicken (*Gallus gallus*), anole lizard (*Anolis carolinensis*) and zebrafish (*Danio rerio*). This reduced number of well-annotated species allowed us to assemble small multi-sequence alignments suitable to carry out short exploratory analysis of our inner ear databases. The list of transcription factors expressed in the inner ear was assembled through bibliographic searches (Table S3). Orthologs finding and coding sequence download were performed through BioMart data mining tool, consulting Ensembl orthologs database. As BioMart downloads one sequence per coding sequence reported for each gene in each species, only the largest coding sequence was kept as criteria to choose that which would best represent the coding sequence of the complete locus. In order to perform an evolutionary analysis on a gene, this must be represented by only one, ortholog in each species. Therefore, two filtering steps were carried on to take away, first those genes for which an ortholog could not be found in one or more of the selected species, and second, those genes that presented orthology relationships of the type one-to-many. Thus a subset of the original genes, suitable to be automatically processed and analyzed, was defined. For deeper analysis of individual genes, coding sequences of all available vertebrate species were downloaded from Ensembl (www.ensembl.org), low quality sequences were excluded and visualization and manual curation of the multiple species alignments were performed in MEGA5 (54), using mainly Genbank Nucleotide (www.ncbi.nlm.nih.gov/genbank) sequence reports. Species tree for each deeply analyzed gene were pruned using ETE Toolkit Python framework (55, 56) from the Ensembl full species tree.

### Sequence Alignment

For the high throughput analysis, sequence alignments were performed with transAlign (57) in order to keep codon-reading frame in DNA sequence. Additionally, a Perl script called batchAlign was implemented to carry out the sequence alignments of multiple genes, in batch (available through Bininda-Emonds at http://www.uni-oldenburg.de/en/biology/systematics-and-evolutionary-biology/). For deeper analysis of individual genes, sequence alignments were performed using Clustal W implemented in MEGA5 software (54).

### Evolutionary Analysis

To carry out the adaptive evolutionary analysis, the modified Model A branch-site test 2 of positive selection (58) was applied in *codeml* program from PAML4 package (59). Two-nested hypotheses were tested for the focal lineage: the alternative hypothesis in which positive selection was allowed only in the selected branch and the null hypothesis where no positive selection was allowed. In the alternative hypothesis three ω values are calculated: ω_0_, ω_1_ and ω_2_ for the codons under negative, null and positive selection, respectively. In the null hypothesis, as no positively selected sites are allowed, only two ω values must be estimated: ω_0_ and ω_1_. The posterior probabilities of the given data to fit the proposed evolutionary model in each hypothesis was obtained by maximum likelihood optimization. Likelihood Ratio Tests (LRTs) were performed from such probability values comparing twice the difference of the log-likelihood probabilities for both hypotheses against a chi-square distribution with 1 degree of freedom. In those cases where the positive selection test rejected the null hypothesis, a-posteriori Bayesian approach to specifically identify the sites under positive selection along the branch of the tree studied was implemented by *codeml* after maximum likelihood estimations were concluded. This Bayes Empirical Bayes (BEB) analysis estimates the posterior probability of each site in the alignment to belong to each of the site classes determined by the model (58).

In the 7 species batch exploratory analysis, as a complete deletion parameter was set, an additional filter was added before the analysis that excluded those genes with such a high proportion of gaps and missing information in their sequence alignments that they could not be processed by PAML.

Hereafter we implemented a method to simplify and lower the computational time required to perform the branch-site positive selection analysis in PAML: we split the optimization of the total number of parameters in this complex model of positive selection test running first the M0 basic model over the species raw topology and obtained the optimized maximum likelihood branch lengths using the Goldman&Yang94 (60) codon based evolutionary model of DNA evolution. Once the branch lengths were estimated, the obtained phylogeny was assigned as the fixed input tree that was used without modifications throughout the branch-site positive selection test. In this way, the branch lengths of the species tree, which comprises an important number of the estimated parameters, are pre-calculated in the M0 run, leaving only the model specific parameters to be optimized during the branch-site test. This approach has shown to be very accurate, and its results highly consistent among reruns of the same data. Our multiPAML in-house Python program mimics this approach, but also automatizes it to make scalable to large quantity of genes. This program also complements the analysis applying a multiple testing correction to the resulting p values through the Benjamini and Hochberg method, commonly knows as False Discovery Rate (FDR) (61).

For deeper analysis of individual genes, in particular those preferentially expressed in OHCs and previously identified to be under positive selection in the mammalian basal lineage, all available vertebrate species were used (Table S8) and other three focal branches were tested: birds basal lineage (Aves), tetrapods basal lineage (Tetrapoda) and tetrapods plus Coelacanth (*Latimeria chalumnae*) lineages.

### In situ hybridization

For *in situ* hybridization (ISH) in cryosections, P8 mice were anesthetized and perfused using 4% paraformaldehyde (PFA) in phosphate-buffered saline (PBS) at 4°C. Cochleas from at least three mice were dissected and fixed in 4% PFA in PBS at 4°C overnight (ON). Cochleas were then treated with a solution containing 0.5mM EDTA pH 8 in PBS for three days at 4°C and embedded in OCT (ThermoFisher Scientific). Blocks were frozen for 1 min in isopentane at -70°C and stored at -80°C. Serial 10-μm-thick sections were cut in a cryostat (Leica 1510S, Germany), collected on Super-Frost Plus slides (FisherScientific, USA), and stored at -80°C until use. Gene coding sequences were cloned into plasmids (pGEMteasy, PROMEGA) through PCR on cDNA using specific primers (Table S9). To synthetize RNA probes, plasmids containing (*Strip2*; *Znf516*; *Ablim2*; *Atp13a1*; *Gcn1l1*; *Synm*; *Lmod3*) were linearized using restriction enzymes, and transcribed using T7 RNA polymerase or SP6 (Roche, Penzberg, Germany) in order to generate digoxigenin (DIG)-labeled sense and antisense riboprobes. After thawing sections, they were postfixed with 4% PFA in PBS for 10 min and then rinsed with PBS for 15 min. Sections were acetylatilated in agitation for 10 minutes (HCl 36%, Acetic anhydride 0.25% v/v, Triethanolamine 1,36% v/v), followed by permeabilization for 30 minutes (1% Triton X-100 in PBS). After washing in PBS, prehybridization was carried out at room temperature for 3 to 4 h in a solution containing 50% formamide, 10% dextran sulfate, 5x Denhardt’s solution, and 250 mg/ml tRNA. Hybridization was performed with 300 ng/ml of the probe in hybridization solution at 72°C for 16h. Thereafter, samples were washed with 0.2% SSC at 72°C for 45 min - 1 hour and then twice for 5 minutes with B1 solution (containing 100 mM NaCl, 0.1% Triton X-100, and 100 mM Tris-HCl pH 7.5). After blocking with 10% normal goat serum in B1 solution for 4 h, the sections were incubated ON with alkaline phosphatase-conjugated anti DIG Fab fragments (Roche, Germany; 1:3,500 in B1). The next day, sections were washed in B1 and incubated in B2 (100 mM NaCl, 50mM MgCl2, 0.1% Tween, and 100 mM Tris-HCl pH 9.5) twice for 10 minutes. The time of the staining reaction began when B2 + NBT-BCIP (Roche, Germany, 0,2% v/v) reactive was added. Samples were checked after 1 h, and every 20 min under a microscope to determine when the detection reaction should be stopped. The reaction was stopped when the signal to noise ratio was at an optimum. Both sense and antisense reactions were stopped at the same time. Finally, samples were coverslipped with a solution containing polyvinyl alcohol and glycerol in Tris 0.2M, pH 8.5. ISH samples were photographed using a microscope Leica DM2500 coupled to a digital camera Leica DFC 7000 T. Images were processed using Adobe Photoshop.

For fluorescent ISH, after blocking with 10% normal goat serum in B1 solution for 4 h, the sections were incubated ON with horseradish peroxidase-conjugated anti DIG Fab fragments (Roche, Germany; 1:500 in B1). The next day, sections were washed in B1 and incubated in Cy3 Amplification Reagent Working Solution (Perkin Elmer) for 30 minutes. Finally, samples were coverslipped with a solution containing Vectashield mounting medium (Vector Laboratories, U.S.A.). Samples were photographed using a Confocal Leica TCS SPE microscope and images were processed using Adobe Photoshop.

### Mutant mice generation

Mouse strains carrying deleted (knockout) alleles were created using a modified CRISPR/Cas9 protocol (Wang et al., 2013). Briefly, sgRNA recognition sequence targeting the *Ablim2* region (5’-agaacaagtacttccacatc AGG-3’, where AGG is the PAM) or the *Strip2* region (5’-atggacgaccccgc ggcacc GGG-3’, where GGG is the PAM) were designed using CRISPR Design Tool (http://crispr.mit.edu/). No potential off-targets were found by searching for matches in the mouse genome (mm10) and allowing for up to two mismatches in the 20 nt sequence preceding the NGG PAM sequence. The T7 promoter was added to the recognition sequence, and the whole sgRNA was generated by a PCR with a reverse primer (5’-aaaagcaccgactcggtgcc-3’) from the pX330 plasmid. The T7-sgRNA product was used as a template for in vitro transcription using the MEGAshortscript T7 kit (ThermoFisher Scientific). The Cas9 mRNA was in vitro transcribed from pMLM3613 plasmid using the mMESSAGE mMACHINE T7 kit (Thermo Fisher Scientific) and polyadenylated using Poly(A) Tailing Kit (Thermo Fisher Scientific #AM1350). Transgenic knockout mice were generated by microinjecting 1 pl of a solution containing Cas9 mRNA (final concentration of 100 ng/ul) and sgRNA (50 ng/ul) in nuclease-free water directly into the cytoplasm of FVB/NJ zygotes and a few hours later transferred to the oviduct of pseudopregnant FVB/NJ females. F0 mice were genotyped by PCR (see genotyping primers in Supp. Table 9) and mutations identified by DNA sequencing. All analyses were performed in littermates resulting of F1 heterozygous mattings (F2). All mouse procedures were performed in agreement with the INGEBI-CONICET Laboratory Animal Welfare and Research Committee.

### Cochlear function tests

To evaluate the integrity of the hearing system we recorded ABRs (Auditory Brainstem Responses) that are sound-evoked potentials generated by neuronal circuits in the ascending auditory pathways (25). We also evaluated the OHCs function through Distortion Product Otoacoustic Emissions (DPOAE) testing, which can be measured from the external auditory canal. When two tones are presented to the normal ear, distortion components at additional frequencies are produced in the hair cell receptor potentials that can drive the OHCs’ electromotility to move the sensory epithelium at the distortion frequencies. The resulting pressure waves from the motion of the sensory epithelium are conducted back to the eardrum to produce DPOAEs, which can be measured in the ear canal (26). ABRs and DPOAEs, were performed on mice anesthetized with xylazine (20 mg/kg, i.p.) and ketamine (100 mg/kg, i.p.) and placed in an acoustically electrically shielded chamber maintained at 30°C. Recordings were performed at postnatal day 60 (P60). Sound stimuli were delivered through a custom acoustic system with two dynamic earphones used as sound sources (CDMG15008–03A; CUI) and an electret condenser microphone (FG-23329-PO7; Knowles) coupled to a probe tube to measure sound pressure near the eardrum (for details, see https://www.masseyeandear.org/research/otolaryngology/investigators/laboratories/eaton-peabody-laboratories/epl-engineering-resources/epl-acoustic-system). Digital stimulus generation and response processing were handled by digital I-O boards from National Instruments driven by custom software written in LabVIEW. For ABRs, needle electrodes were placed into the skin at the dorsal midline close to the neural crest and pinna with a ground electrode near the tail. ABR potentials were evoked with 5 ms tone pips (0.5 ms rise-fall, with a cos^2^ envelope, at 40/s) delivered to the eardrum at log-spaced frequencies from 5.6 to 45.25 kHz. The response was amplified 10,000X with a 0.3–3 kHz passband. Sound level was raised in 5 dB steps from 10 to 80 dB sound pressure level (SPL). At each level, 1024 responses were averaged with stimulus polarity alternated. The DPOAEs in response to two primary tones of frequency f1 and f2 were recorded at (2 × f1)-f2, with f2/f1=1.2, and the f2 level 10 dB lower than the f1 level. Ear-canal sound pressure was amplified and digitally sampled at 4 µs intervals. DPOAE threshold was defined as the lowest f2 level in which the signal to noise floor ratio is >1.

### Cochlear processing and immunostaining

Cochleae were extracted, perfused intra-labyrinthly with 4% PFA in PBS, post-fixed with 4% PFA ON and decalcified in 0,12M EDTA for five days. Cochlear tissues were then microdissected and permeabilized by freeze/thawing in 30% sucrose (for CtBP2/GluA2 immunostaining) or directly blocked (for Ablim/prestin immunostaining). The microdissected pieces were blocked in 5% normal goat serum (for CtBP2/GluA2 immunostaining) or 5% normal donkey serum (for Ablim/prestin immunostaining) with 1% Triton X-100 in PBS for 1 h, followed by incubation in primary antibodies (diluted in blocking buffer) at 37°C for 16 h (for CtBP2/GluA2 immunostaining) or 4°C for 16 h (for Ablim/prestin immunostaining). The primary antibodies used in this study were: 1) rabbit anti-ablim antibody (rabbit polyclonal anti Ablim2, abcam ab100926;1:200) 2) goat anti-prestin antibody (Santa cruz biotechnology inc. sc22692;1:700) 3) anti-C-terminal binding protein 2 (mouse anti-CtBP2 IgG1; BD Biosciences, San Jose, CA; 1:200) and 4) anti-glutamate receptor 2 (mouse anti-GluA2 IgG2a; Millipore, Billerica, MA; 1:2000). For F-actin detection we used TRITC-Phalloidin (Sigma). Tissues were then incubated with the appropriate Alexa Fluor-conjugated fluorescent secondary antibodies (Invitrogen, Carlsbad, CA; 1:1000 in blocking buffer) for 2 h at room temperature. Finally, tissues were mounted on microscope slides in FluorSave mounting media (Millipore, Billerica, MA). Confocal z-stacks (0.2 µm step size) of the medial region from each cochlea were taken using a Leica TCS SPE microscope equipped with 63X (1.5X digital zoom) oil-immersion lens.

### Western blot analyses

To analyze Strip2 expression in mutated and wild type mice we performed western blot assays using heart and liver protein extracts prepared in a protein extraction buffer [50 mM Tris-HCl pH 7.5, 2 mM EDTA; 1% Triton X100; 150 mM NaCl; 0.05% SDS; Halt™ Protease and Phosphatase Inhibitor Cocktail (100X) (Thermo Scientific 78440)]. Then we added 150 mM NaCL, 0,2 % glycerol, 2% bromophenol blue and β-Mercaptoethanol to 10% to the protein extract and heated them at 100°C (5min). Samples were then separated on 12% SDS-Polyacrylamide gels (prepared with Acrilamide and N,N′-Methylenebisacrilamide 30%, Invitrogen) and transferred using a wet transfer to nitrocellulose membranes (BIO-RAD). Membranes were blocked in 5% (w/v) nonfat dry milk, 0.05% v/v Tween 20 in TBS (milk/1xTBS-T) for 1 hour. After blocking, membranes were incubated overnight at 4°C in blocking solution containing the primary antibody. The membrane was probed with a polyclonal anti-human STRIP2 antibody produced in rabbit (HPA019657 Sigma) at a dilution of 1:500 according to the manufacturer’s protocol. After washing 3 times in TBS containing 0.05% v/v Tween 20, blots were incubated with the appropriate secondary antibody donkey anti-rabbit HRP conjugate (1:2000, Fisher Scientific) for 3 hours at room temperature. Polyclonal anti-human beta Tubulin antibody produced in rabbit (Abcam Ab6046) was used as loading control at a dilution 1:500. Proteins were visualized using ECL detection (Cell Signaling Technology SignalFire™ ECL Reagent #6883) and by exposing on the GeneGnomeXRQ (Syngene).

## Author Contributions

LFF designed and supervised the project. FP, MS, ARC, VC, MEGC and LFF conducted experiments and data analysis. LFF wrote the manuscript. MR, ABE and MEGC discussed experiments, provided reagents and materials and edited the manuscript. All authors edited and approved the final version of this report.

## Acknowledgements

We thank Marta Treimun for superb technical assistance with mutant mice generation.

---

**Supplementary Figure 1. Strip2 is expressed in the cochlear nuclei in adult mice.**

(a) *Strip2* expression analysis through ISH performed by the Allen Brain Institute (www.alleninstitute.org) showing the expression at the cochlear nuclei. (b) Higher magnification showing dorsal and ventral cochlear nuclei. ISH experiments confirmed the expression at the dorsal CN in wild type (c) and *Strip 2* mutant mice (d).

